# Autophagy alterations in white and brown adipose tissues of mice exercised under different training protocols

**DOI:** 10.1101/2022.09.05.505110

**Authors:** Isaac Tamargo-Gómez, Manuel Fernández-Sanjurjo, Helena Codina-Martínez, Cristina Tomás-Zapico, Eduardo Iglesias-Gutiérrez, Álvaro F. Fernández, Benjamín Fernández-García

## Abstract

Autophagy is a conserved catabolic process that promotes cellular homeostasis and health. Although exercise is a well-established inducer of this pathway, little is known about the effects of different types of training protocols on the autophagy levels of tissues that are tightly linked to the obesity pandemic (like brown adipose tissue) but not easily accessible in humans. Here, we take advantage of animal models to assess the effects of short- and long-term resistance and endurance training in both white and brown adipose tissue, reporting distinct alterations on autophagy proteins LC3B and p62. For instance, both short-term endurance and resistance training protocols increased the levels of these proteins in white adipose tissue before this similarity diverges during long training, while autophagy regulation appears to be far more complex in brown adipose tissue. Additionally, we also analyzed the repercussion of these interventions in fat tissues of mice lacking autophagy protease ATG4B, further assessing the impact of exercise in these dynamic, regulatory organs (which are specialized in energy storage) when autophagy is limited. In this regard, only resistance training could slightly increase the presence of lipidated LC3B, while p62 levels increased in white adipose tissue after short-term training but decreased in brown adipose tissue after long-term training. Altogether, our study suggests an intricated regulation of exercise-induced autophagy in adipose tissues that is dependent on the training protocol and the autophagy competence of the organism.

## INTRODUCTION

The local and systemic benefits of exercise training have been widely described in the literature^1^, not only for sports performance but also to prevent and treat several chronic pathologies, including metabolic disorders^2^. Thus, exercise has emerged as a powerful and safe lifestyle intervention to promote health throughout the lifespan, including elderly^3^. Nine major hallmarks have been defined to characterize aging^4^, with exercise being able to counteract several of them^5,6^. Among these hallmarks of aging, as characterized by López-Otín and colleagues, is the loss of autophagy, one of the cell’s main proteolytic systems. In fact, its decline has been described during aging in different animal models^7,8^, while its activation has been linked to increased lifespan and healthspan in flies, worms, and mammals^9–11^. Endurance exercise is a well-known physiological inducer of autophagy in skeletal muscle^12^ and we have described that this mechanism is particularly relevant for resistance gain and exercise-induced adaptations in brain^13^. However, the effects of exercise training on the autophagic response of white and brown adipose tissues (WAT and BAT) remain unclear. This is relevant, as autophagy and adipose tissues play an important role in the maintenance of the metabolic balance of the organism and for the prevention of metabolic, cardiovascular, and cognitive diseases^14–17^. Furthermore, it has been described that endurance exercise induces WAT “browning”^18^, a process that has been linked with a diminished risk of metabolic diseases^19^ and in which autophagy plays an important role^20^. Nevertheless, very little is known about the effect of resistance exercise on autophagy in adipose tissues.

Here, we describe how different exercise models (resistance and endurance, short- and long-term) modify the levels of autophagy markers in WAT and BATs. Moreover, we use autophagy-deficient mice to describe how the impact of training is affected when this process is hampered, as it is observed during aging.

## MATERIAL AND METHODS

### Animals

8-week-old male mice with mixed background C57BL6/129Sv, deficient in *Atg4b*^21^ (KO; n = 36) and their corresponding wild-type controls (WT; n = 36), were used. Eight mice were housed per cage with food and water *ad libitum*. Mice were maintained on a 12 h light/dark cycle (onset at 8:00 AM) and under controlled temperature (22 ± 2 ºC). Two training periods were conducted: 2-week intervention (48 animals, n = 24 per genotype) and 14-week intervention (24 animals, n = 12 per genotype), which is the minimum period in human training studies. Mice were randomly and equally distributed into 3 groups for each intervention: sedentary, resistance training, and endurance training. All procedures were conducted during the light portion of the cycle, between 8:30 and 11 AM, at the Animal Facility of Universidad de Oviedo, and were performed in accordance with institutional guidelines approved by the Committee on Animal Experimentation of Universidad de Oviedo (PROAE 30/2016).

### Physical training methods

A commercial treadmill for rats (Panlab 8700, Barcelona, Spain) and an own-manufactured vertical ladder were used for training. The ladder was built with 30 steel wire steps of 1.5 mm diameter, separated by 15 mm. A 20 × 20 cm resting area was placed on the top of the ladder. The slope of the ladder could vary from 70° to -80° with the horizontal plane. Two lanes were delimited to prevent non-linear climbing.

#### Acclimation period (2 weeks)

To favor mice acclimatization, all animals stayed 20 min in each working station (treadmill, ladder, and clean cage), 5 days/week in groups of four. The first week, mice were placed on the treadmill without movement and in the resting area at the top of the ladder. The following week, animals walked on the moving treadmill belt (10 cm/s, for 20 minutes) and they were shown how to climb the ladder, from the 5^th^ top step to the resting area, with 2 minutes of rest intervals, for 20 minutes.

#### Training period (2 weeks or 14 weeks)

The mice trained in groups of four, with no rejection to training. No aversive stimuli were used. Progressive and humanized training protocols were performed^22^. For endurance training, the duration of the sessions, the maximal speed, and the slope of the treadmill were gradually increased along the training period. For resistance training, the number of series, the number of steps per series, and maximal weight were progressively increased. In each training session, the starting speed for endurance training was 50% of the maximal running speed of the weakest mouse. For resistance training, the initial weight was 5% of the maximal weight in the performance test. Control mice explored freely in a new cage while the exercise groups were training in the same room.

### Immunoblotting

Mice were sacrificed 48 hours after the last exercise bout by CO_2_ inhalation. Perigonadal WAT and interscapular BAT were extracted and preserved at -80 ºC. Then, they were homogenized in an extraction buffer consisting of 20 mM HEPES, pH 7.4, 100 mM NaCl, 50 mM NaF, 5 mM EDTA, 1% Triton X-100, 1 mM sodium orthovanadate, 1 mM pyrophosphate, and Complete protease inhibitor cocktail (Hoffmann-La Roche, Basel, Switzerland). Then, samples were centrifuged at 12,000 *g*, at 4 ºC, for 10 minutes, and the supernatant was collected and preserved at -80 ºC until further use. Protein quantification was determined by the bicinchoninic acid technique (Pierce BCA Protein Assay kit; Pierce Biothecnology Inc., Waltham, MA, USA). Total protein (20 μg) was electrophoresed on 15% SDS-polyacrylamide gel and transferred to PVDF (Millipore, Burlington, MA, USA) membranes, which were then blocked in TBS-T (Tris-buffered saline with 5% BSA and 1% Tween-20). Next, membranes were incubated with rabbit anti-LC3B (NB600-1384; Novus Biologicals, Littleton, CO, USA) and mouse anti-SQSTM1/p62 clone 2C11 (H00008878-M01; Abnova, Taipei, Taiwan) antibodies diluted in TBS-T. Then, membranes were incubated with corresponding secondary antibodies (Santa Cruz Biotechnologies, Dallas, TX, USA), conjugated with horseradish peroxidase, during 1 hour at RT. Detection was developed with Luminata™ Forte (Millipore, Burlington, MA, USA) and images acquired using the software VisionWorksLs (UVP, Upland, CA, USA). Anti-actin conjugated with horseradish peroxidase (sc-47778; Santa Cruz Biotechnologies, Dallas, TX, USA) was used as a loading control.

### Statistical analysis

Data are presented as means ± SEM and represented as column scatter plots. Normality of the variables was tested by means of the Shapiro-Wilk test. Comparison between the three experimental groups was performed using a one-way ANOVA test with Tukey’s post-hoc test. Graph Pad Prism 7 was used for statistical analysis (Graph Pad Software, La Jolla, CA, USA). A critical value for the significance of *P*< 0.05 was considered throughout the study.

## RESULTS

### Training alters the levels of autophagy markers in adipose tissue

After describing the different effects of endurance and resistance training on the levels of autophagy in the brain of rested and exercised mice^13^, we decided to address the status of autophagy markers in the adipose tissue after *2 weeks* or *14 weeks* of the aforementioned training protocols. For this purpose, we performed immunoblotting analyses to detect autophagy markers LC3B (as the accumulation of its lipidated form, LC3B-II, is associated with increased autophagy) and SQSTM1/p62 (an autophagy substrate)^23^ in both gonadal white adipose tissue (WAT) and interscapular brown adipose tissue (BAT). Interestingly, both endurance and resistance training protocols increased the levels of these autophagy markers in WAT after just *2 weeks* of exercise (Figure 1A and B). In the case of endurance training, it also resulted in an increased ratio of the lipidated form of LC3B (LC3B-II) over the unconjugated one (LC3B-I), which could hint at a possible accumulation of autophagic vesicles within these cells. On the contrary, exercising the mice for *14 weeks* distinctly affected the expression of these proteins (Figure 1C and D). For instance, endurance training significantly reduced the levels of LC3B-II while augmenting the presence of p62 in WAT. Meanwhile, a slight reduction in the quantity of LC3B-II after resistance training was not accompanied by an accumulation of p62, as its levels remained comparable to those from rested mice. Thus, while short exercise protocols seem to induce the accretion of autophagy markers in WAT, the extension of these interventions results in the differential regulation of their expression depending on the type of exercise. However, these changes are not only exercise- or extension-dependent, but also tissue-dependent. For instance, *2 weeks* of endurance training also caused LC3B-II accumulation in BAT, accompanied by a p62 reduction, in opposition to what we observed in WAT (Figure 1E and F). As for resistance training, this short exercise protocol resulted in a significant decrease of both p62 and LC3B-II when these samples are compared to those from control animals. Intriguingly, the 14-week-long exercise protocols had comparable effects on the levels of autophagy markers in BAT (Figure 1G and H), as both endurance and resistance training increased LC3B-II/LC3B-I ratio while the amounts of p62 remain unaltered. Taken together, these results confirm that different training protocols can distinctly alter the status of autophagy markers in adipose tissues.

**Figure 1.**
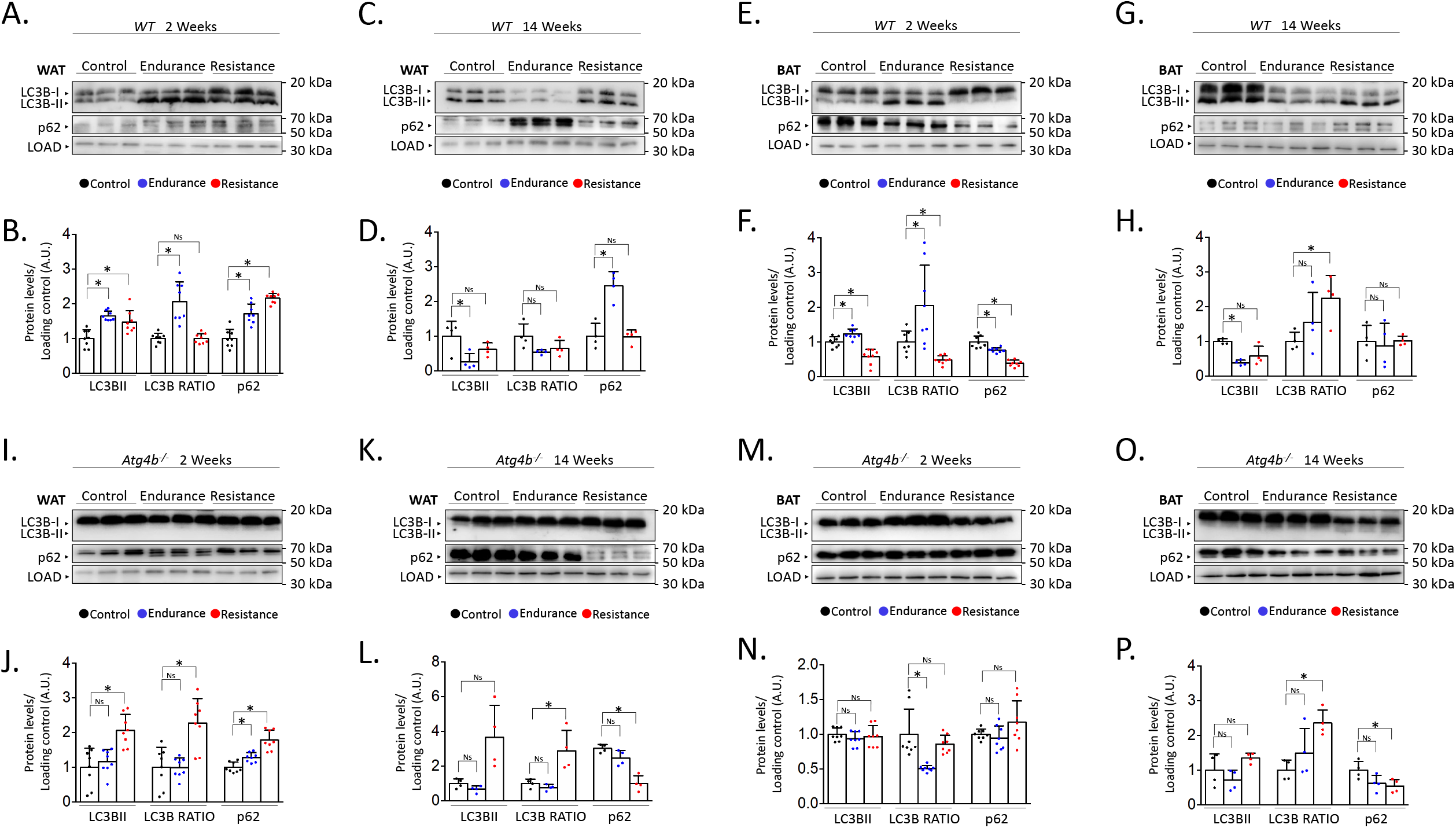
Alterations on autophagy markers in white and brown adipose tissues after endurance or resistance training. Western blot analysis of autophagy markers LC3B and p62 in white adipose tissue from wild-type mice that were trained for 2 weeks. **(B)** Quantification of the data from (A). **(C)** Western blot analysis of autophagy markers LC3B and p62 in white adipose tissue from wild-type mice that were trained for 14 weeks. **(D)** Quantification of the data from (C). **(E)** Western blot analysis of autophagy markers LC3B and p62 in brown adipose tissue from wild-type mice that were trained for 2 weeks. **(F)** Quantification of the data from (E). **(G)** Western blot analysis of autophagy markers LC3B and p62 in brown adipose tissue from wild-type mice that were trained for 14 weeks. **(H)** Quantification of the data from (G). **(I)** Western blot analysis of autophagy markers LC3B and p62 in white adipose tissue from *Atg4b*^*-/-*^ mice that were trained for 2 weeks. **(J)** Quantification of the data from (I). **(K)** Western blot analysis of autophagy markers LC3B and p62 in white adipose tissue from *Atg4b*^*-/-*^ mice that were trained for 14 weeks. **(L)** Quantification of the data from (K). **(M)** Western blot analysis of autophagy markers LC3B and p62 in brown adipose tissue from *Atg4b*^*-/-*^ mice that were trained for 2 weeks. **(N)** Quantification of the data from (M). **(O)** Western blot analysis of autophagy markers LC3B and p62 in brown adipose tissue from *Atg4b*^*-/-*^ mice that were trained for 14 weeks. **(P)** Quantification of the data from (O). N = 8 mice per genotype in 2-weeks-long training protocols; N = 4 mice per genotype in 14-weeks-long training protocols. \**P* < 0.05, one-way ANOVA test followed by Tukey’s post-hoc test. WAT: White adipose tissue; BAT: Brown adipose tissue.

### Distinct effects of training in autophagy-deficient adipose tissue

Given the observed results in adipose tissue from trained WT mice, we were prompted to analyze the effect of training in the context of autophagy deficiency. Thus, we trained mice lacking cysteine protease ATG4B, a murine model that shows attenuated autophagic response^21^ and increased susceptibility to different experimentally-induced pathologies^24– 27^. Even more, we have previously described that these animals show worse adaptive responses to exercise, also presenting alterations in the levels of autophagy markers in different brain areas^13^. As it is shown in Figures 1I and 1J, both endurance and resistance training protocols induced a slight accumulation of p62 in the WAT of exercised animals after two weeks in comparison to their rested *Atg4b*-deficient littermates. However, only resistance exercise was able to increase the presence of LC3B-II in this tissue. ATG4B-deficiency significantly disrupts LC3B-II formation in these animals^21^, which suggests that the detection of the lipidated form of LC3B after resistance training may be due to a compensatory upregulation on the activity of the other ATG4 proteases (ATG4A, ATG4C and/or ATG4D). When the protocol is extended up to *14 weeks*, mice exercised under the endurance training protocol showed protein levels that were comparable to those from control *Atg4b*^*-/-*^ animals (Figure 1K and L). However, WAT from mice that trained resistance still showed increased LC3B-II levels and increased LC3B-II/LC3B-I ratio, even though p62 was now significantly reduced. Curiously, 2-week training protocols had little to no effect on the levels of autophagy markers in BAT from *Atg4b*-deficient mice (Figure 1M and N), as all three groups exhibited similar levels of LC3B-II and p62. In these animals, the LC3B-II/LC3B-I ratio was decreased after endurance training, though this was due to the accumulation of LC3B-I in this group. As for the 14-week-long training intervention, mice that trained resistance displayed once again a statistically significant reduction on p62 with an increase in LC3B-II/LC3-I ratio, similar to what we observed in WAT (Figure 1O and P). Protein levels were not significantly different after endurance training, though we could detect a partial trend towards p62 decrease. Collectively, these data confirm that autophagy markers are also altered upon exercise in the adipose tissues of mice lacking ATG4B protease.

Last, we decided to compare protein levels of LC3B and p62 between WT and *Atg4b*^*-/-*^ animals within the same group. This approach allowed us to 1) assess possible alterations in these autophagy markers in basal conditions due to the absence of ATG4B and 2) measure how the effect of training is affected when autophagy is hampered. In WAT, rested knock-out mice showed reduced LC3B-II levels and LC3B-II/LC3B-I ratio, as well as an accumulation of p62 (Figure 2A-H) when compared to WT animals that were not trained either. These results further confirm our previous observations on the status of autophagy markers in the white adipose tissue of *Atg4b*-deficient mice^27^. All training protocols (endurance and resistance, short and long) failed to rescue LC3B or LC3B-II/LC3B-I ratio to the levels displayed by the corresponding WT groups in this tissue, highlighting the crucial role of ATG4B in LC3-II formation. As for p62, *Atg4b*^*-/-*^ mice still showed increased levels of this marker after the 14-week-long endurance training and the 2-week resistance training. When we compared protein levels in BAT, we found out that *Atg4b*-deficient mice also showed a dramatic reduction of LC3B-II and also in LC3B-II/LC3B-I ratio. However, contrarily to what we observed in WAT, p62 levels were at first reduced in younger knock-out mice but already augmented after 14 weeks (Figure 2I-P), suggesting a probable accumulation in this tissue with ageing, and matching the phenotype we describe in WAT. Once again, none of the training protocols succeeded in rescuing LC3B-II levels or LC3B-II/LC3B-I ratio to what is displayed by those observed in trained WT mice. Interestingly, while short training periods had either a null (endurance) or accumulative effect (resistance) on the levels of p62, prolonged training resulted in a significant reduction of this protein in brown adipose tissue of knock-out mice. Altogether, these findings further demonstrate that exercise distinctly alters autophagy markers depending on the tissue, the protocol, and the presence/absence of ATG4B (Figure 3).

**Figure 2.**
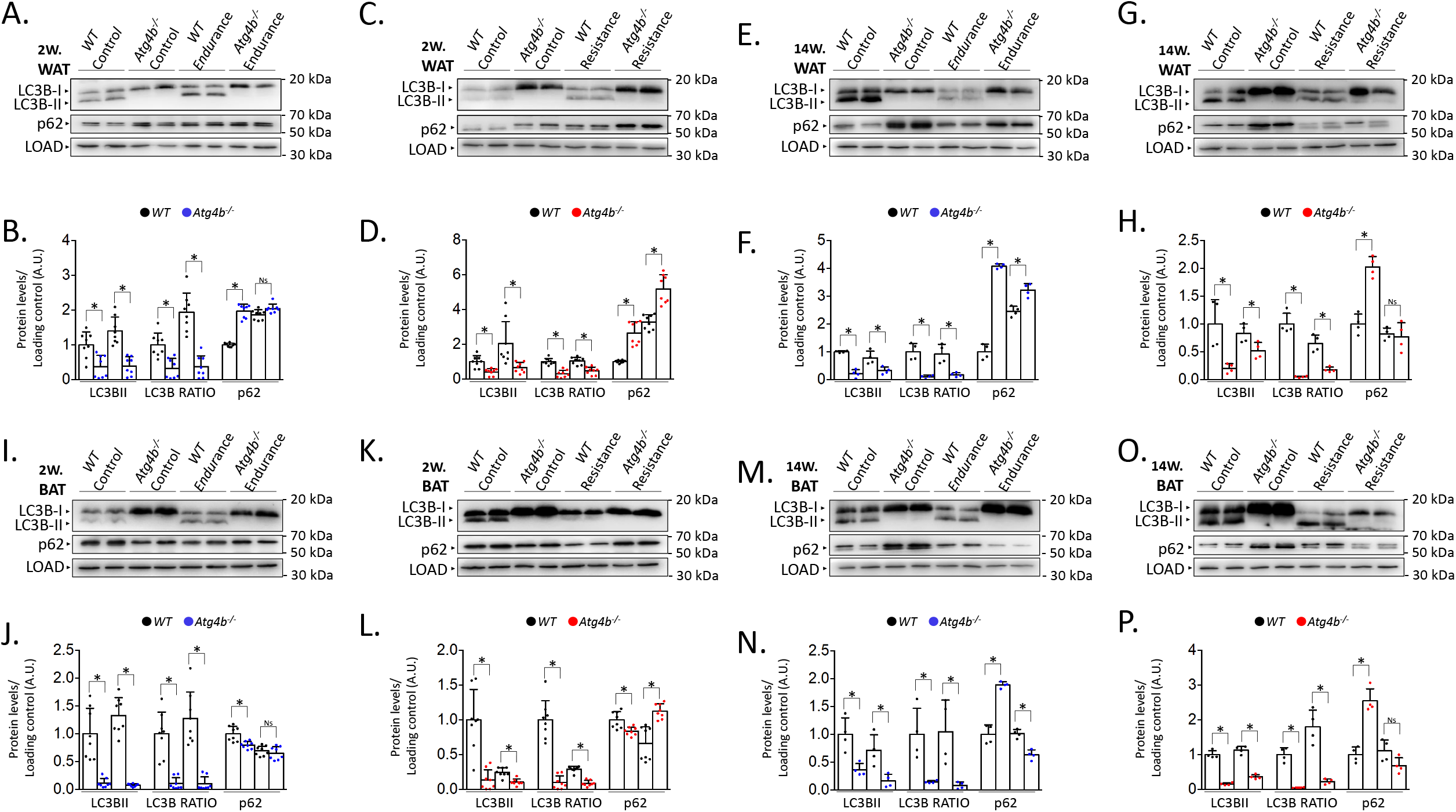
Comparison of the levels of autophagy markers in white and brown adipose tissue from wild-type and *Atg4b*^*-/-*^ mice. **(A)** Western blot analysis of autophagy markers LC3B and p62 in white adipose tissue from wild-type or *Atg4b*^*-/-*^ mice that have either rested or exercised for 2 weeks following an endurance training protocol. **(B)** Quantification of the data from (A). **(C)** Western blot analysis of autophagy markers LC3B and p62 in white adipose tissue from wild-type or *Atg4b*^*-/-*^ mice that have either rested or exercised for 2 weeks following a resistance training protocol. **(D)** Quantification of the data from (C). **(E)** Western blot analysis of autophagy markers LC3B and p62 in white adipose tissue from wild-type or *Atg4b*^*-/-*^ mice that have either rested or exercised for 14 weeks following an endurance training protocol. **(F)** Quantification of the data from (E). **(G)** Western blot analysis of autophagy markers LC3B and p62 in white adipose tissue from wild-type or *Atg4b*^*-/-*^ mice that have either rested or exercised for 14 weeks following a resistance training protocol. **(H)** Quantification of the data from (G). **(I)** Western blot analysis of autophagy markers LC3B and p62 in brown adipose tissue from wild-type or *Atg4b*^*-/-*^ mice that have either rested or exercised for 2 weeks following an endurance training protocol. **(J)** Quantification of the data from (I). **(K)** Western blot analysis of autophagy markers LC3B and p62 in brown adipose tissue from wild-type or *Atg4b*^*-/-*^ mice that have either rested or exercised for 2 weeks following a resistance training protocol. **(L)** Quantification of the data from (K). **(M)** Western blot analysis of autophagy markers LC3B and p62 in brown adipose tissue from wild-type or *Atg4b*^*-/-*^ mice that have either rested or exercised for 14 weeks following an endurance training protocol. **(N)** Quantification of the data from (M). **(O)** Western blot analysis of autophagy markers LC3B and p62 in brown adipose tissue from wild-type or *Atg4b*^*-/-*^ mice that have either rested or exercised for 14 weeks following a resistance training protocol. **(P)** Quantification of the data from (O). N = 8 mice per genotype in 2-weeks-long training protocols; N = 4 mice per genotype in 14-weeks-long training protocols. \**P* < 0.05, one-way ANOVA test followed by Tukey’s post-hoc test. WAT: White adipose tissue; BAT: Brown adipose tissue.

**Figure 3.**
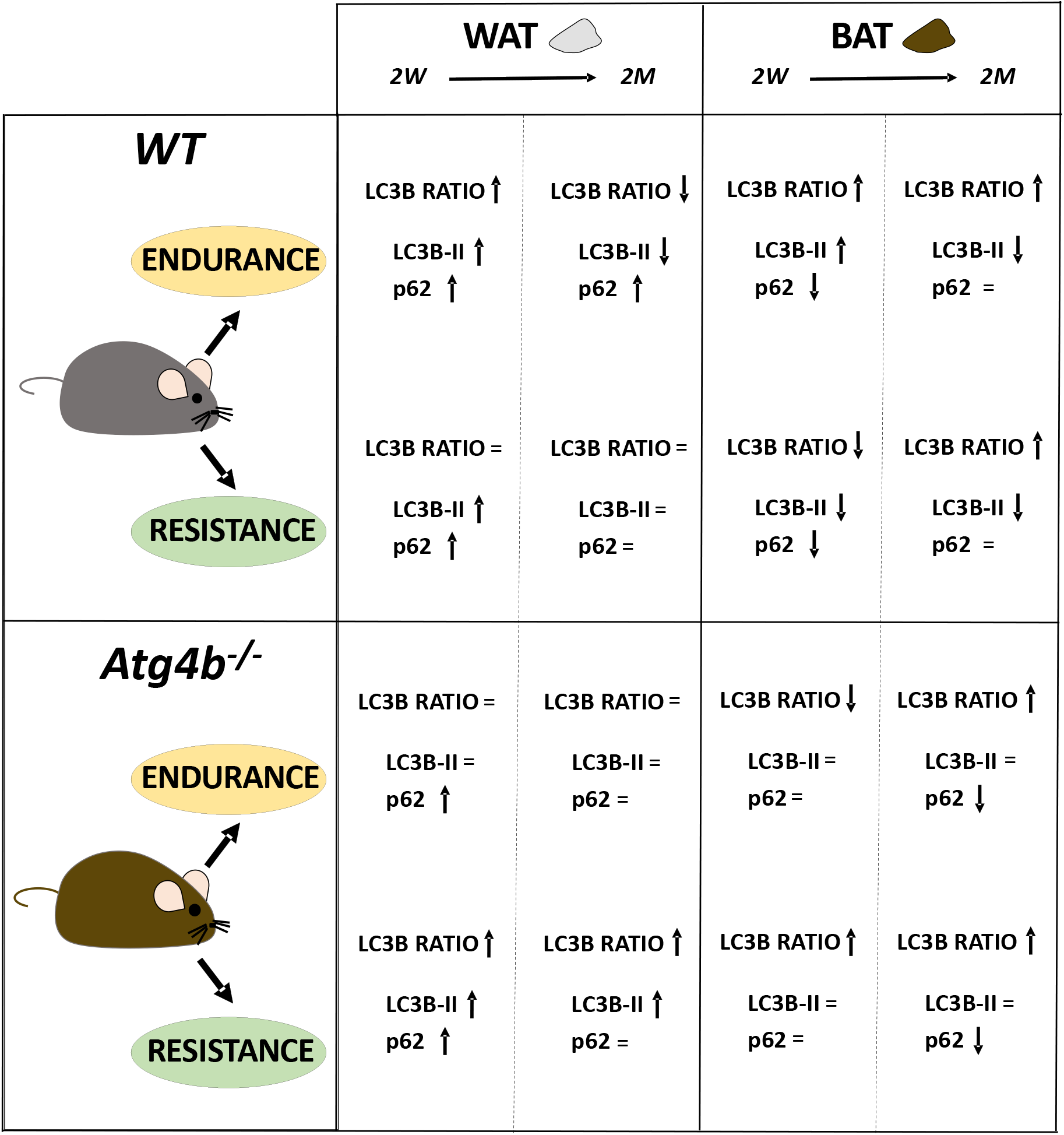
The complex regulation of autophagy in adipose tissues after endurance or resistance training. Diagram depicting the alterations on the levels of autophagy markers in white and brown adipose tissues from wild-type and *Atg4b*^*-/-*^ mice as a result of different training protocols.

## DISCUSSION

Our knowledge on the link between autophagy and exercise has expanded since it was first reported^12^, yet we are still far from completely understand the role of this cellular process in training adaptation of different organs and tissues. Even though recent articles have described how endurance training affects autophagy in white adipose tissue of different animal models^12,28–30^, this is to the best of our knowledge the first study to compare and analyze different types of exercise (endurance and resistance), lengths of training protocols (*2 weeks* and *14 weeks*) and fat tissues (WAT and BAT) in both WT and autophagy-deficient mice. In this regard, the use of animal models with partial autophagy capacity facilitates the investigation of how autophagy decline, which has been already described during aging^9^ and disease^31^, can hamper the beneficial effects of exercise on the elderly. Moreover, animal models allow the study of organs that are not easily accessible like brown adipose tissue, which is tightly linked to the obesity pandemic. In this regard, assessing the relevance of exercise-induced autophagy in adipocytes is important as training is considered one of the main strategies to counteract the consequences of hypercaloric diets and sedentarism. Furthermore, the observed dynamic, complex regulation of autophagy in the adipose tissues could have been favoured by what is known as the “thrifty genotype”, a hypothesis that aims to explain the persistence of genes that mediate nutrient storage throughout human evolution^32^.

Still, additional analyses with complementary techniques would further expand our knowledge on autophagy, exercise, and adipose tissue homeostasis. The effects of training on WAT and BAT are already being established^33,34^, as well as the association between WAT reduction, BAT increment, and decreased cardiometabolic risk^18,19,35^. At the same time, it is well-known that autophagy plays an important role in the homeostasis of both WAT and BAT^20^, and its dysregulation in WAT leads to metabolic disorders^36^. An interesting question arises when considering that exercise is a direct, physiological inducer of autophagy, hinting at the possibility that the autophagic route is mediating the beneficial effects of training in the function of the adipose tissue. In this regard, our work shows that exercise, similar to what has been described in other tissues, also alters autophagy in adipose tissue in a multifactorial-dependent way. However, it is important to acknowledge that, here, we characterize the basal protein levels of LC3B-II and p62 by western blot analysis. This is a good initial approach to get valuable information about the autophagy status in a given system. However, future studies should address the use of autophagy inhibitors (such as leupeptin) or activators to fully characterize the autophagy flux *in vivo* conditions^37^.

In conclusion, this study shows for the first time a distinct effect of different specific training protocols in the levels of autophagy markers in WAT and BAT tissues. Moreover, using ATG4B-deficient mice with a hampered autophagic response, we observed that an alteration in the correct development of autophagy cannot be fully rescued by training protocols, which is essential when trying to understand if exercise is beneficial during aging when autophagy declines.

## ACKNOWLEDGEMENTS

We thank Dr. Guillermo Mariño for shared reagents and helpful comments. This work was supported by Ministerio de Economía y Competitividad under grant DEP2012-39262 to EIG and DEP2015-69980-P to BFG. IT-G work is supported by Ministerio de Educación, Cultura y Deporte with an FPU grant for PhD studies. MF-S acknowledges receiving financial support from Fundación para la Investigación y la Innovación Biosanitaria del Principado de Asturias (FINBA), “Convocatoria de contratos predoctorales para grupos de investigación del ISPA”.

## CONFLICT OF INTEREST

The authors declare no conflicts of interest.

## Notes

### Competing Interest Statement

The authors have declared no competing interest.

### Summary of Updates

We have noticed that p62 blots in Figures 2M and 2O (which, in any case, show the same pattern of protein levels) were interchanged. While this mistake does not change the results or interpretations of the study, we have corrected it in this version.

